# Brightness demixing for simultaneous multi-target imaging in 3D single-molecule localization microscopy

**DOI:** 10.1101/2025.03.02.639924

**Authors:** Laurent Le, Surabhi K. Sreenivas, Emmanuel Fort, Sandrine Lévêque-Fort

## Abstract

Single-Molecule Localization Microscopy (SMLM) has revolutionized high-resolution imaging, but the simultaneous detection of multiple fluorophores traditionally relies on spectral-based separation, which is inherently constrained by spectral overlap. Here, we introduce Brightness Demixing, a novel method for fluorophore discrimination that exploits brightness, which directly depends on the fluorophores extinction coefficient and quantum yield. By oversampling blinking events, we precisely quantify photon flux as a proxy for brightness, enabling robust differentiation of fluorophores independent of their spectral properties, without requiring additional spectral separation. Brightness Demixing operates within a single detection channel, eliminating the need for additional spectral filters or cameras. We demonstrate this approach with simultaneous two- and three-target imaging in both 2D and 3D configurations. By maintaining single-wavelength excitation and minimizing chromatic aberrations, this method significantly enhances multiplexing in SMLM while remaining fully compatible with existing setups. Brightness Demixing thus offers a simple yet powerful approach for expanding multi-target imaging capabilities in super-resolution microscopy.

## 1 Introduction

Single-Molecule Localization Microscopy (SMLM) has redefined biological imaging by providing nanoscale resolution, far beyond the diffraction limit. A key challenge in SMLM is the simultaneous imaging of multiple molecular targets, which is essential for deciphering biomolecular organization, protein assemblies, and dynamic interactions within cells [1, 2]. Multi-target imaging plays a crucial role in various applications, particularly in spatial proteomics, where it enables researchers to map complex cellular architectures with nanometric precision [3]. However, conventional multiplexing strategies rely primarily on spectral-based separation, which imposes significant limitations on the number of fluorophores that can be simultaneously detected and requires careful chromatic aberration correction.

To overcome these challenges, various multi-color SMLM approaches have been developed. The most common strategy involves combining multiple laser lines and fluorophores with distinct spectral properties [4–17], as demonstrated in techniques such as PALM (Photo-Activated Localization Microscopy) [18–20], STORM (Stochastic Optical Reconstruction Microscopy) [21, 22], and DNAPAINT [23, 24]. While effective, these methods require multiple excitation wavelengths, dedicated detection channels, and precise chromatic corrections to mitigate optical distortions. In some cases, sequential imaging approaches in STORM with successive labeling [25, 26] and DNA-PAINT with the addition of an orthogonal imager [24, 27, 28] can be used to overcome spectral limitations, but at the cost of dramatically increased acquisition times.

Alternative strategies have been proposed to enable simultaneous multi-target imaging with minimal spectral separation. Spectral demixing (SD) is one such approach, which involves the use of dichroic mirrors to split fluorescence signals into separate detection channels, allowing fluorophores with emission maxima shifted by about 20 to 30 nm to be distinguished based on their relative intensities in each channel [29–34]. However, spectral demixing reduces the signal-to-noise ratio (SNR) and requires complex post-processing to associate localizations between channels, often leading to the rejection of a substantial fraction of detected molecules (up to 40% [34]). An alternative solution is high-resolution spectral imaging, which introduces dispersive elements in the detection path to achieve finer spectral discrimination [35–37]. While this approach minimizes cross-talk and allows simultaneous multi-target imaging with a single excitation wavelength, it comes at the cost of increased instrumentation complexity and the need for an additional detection channel.

Beyond spectral-based techniques, fluorescence lifetime-based approaches have also been explored for multi-target imaging [38–40]. By leveraging differences in fluorophore lifetime, these methods provide an alternative contrast mechanism that is independent of spectral properties. However, lifetime-based imaging requires pulsed laser excitation and picosecond-resolved detection, making it more technically demanding and less accessible for widespread adoption in standard SMLM setups.

In this paper, we leverage an unexploited intrinsic property of fluorophores, their brightness, to achieve fluorophore differentiation. Brightness is determined by the number of photons emitted per unit time. While this quantification is uniquely feasible in SMLM at the single-molecule level, as fluorophores are detected individually, it requires specific acquisition conditions. Standard SMLM detection is primarily optimized to measure the total intensity of individual blinking events for localization, without explicitly resolving the photon flux. Typically, exposure times are set to match the average blinking duration of emitters. To extract brightness, fluorophores must remain in the ON state throughout the entire camera exposure time, meaning that brightness quantification necessitates an acquisition regime where individual blinking events are oversampled.

Fluorescence flux has already been used in SMLM but mainly to extract spatial localization from structured light excitation geometries [41–45]. We demonstrate here that brightness can also be used for multi-target imaging, thus adding a new identification parameter at the single-molecule level. Brightness demixing is therefore a more straightforward alternative strategy to apply for any optical setup since it does not require a second channel in the detection system. This approach benefits of a single wavelength excitation which enables the observation of spectrally overlapping fluorophores minimizing chromatic aberration. Brightness demixing enables to image several targets simultaneously, which reduces the experiment time to a single acquisition. In this article, we performed brightness demixing in DNA-PAINT to identify up to three different proteins in 2D imaging. We further extend this approach to 3D imaging using two distinct axial localization strategies: PSF engineering and supercritical emission.

## 2 Principle and implementation

The brightness *β* of a fluorophore, used to characterize its fluorescence efficiency, is defined as *β* = *σ*(*ν*_*exc*_)*ϕ* where *σ*(*ν*_*exc*_) is the molecular extinction cross-section at the excitation frequency *ν*_*exc*_, and *ϕ* is the quantum yield. The molecular extinction cross-section represents the probability of photon absorption and is an intrinsic characteristic of the molecule. In contrast, the quantum yield corresponds to the probability that an absorbed photon leads to fluorescence photon rather than non-radiative relaxation. Note that the brightness is also expressed for a mole *β* = *ϵ*(*ν*_*exc*_)*ϕ, ϵ*(*ν*_*exc*_) being the molar decadic extinction coefficient.

The radiative part of *ϕ* depends on the local electromagnetic environment [46]. In biological samples, this electromagnetic environment can be considered homogeneous throughout the sample. Small variations can occur in the vicinity of the coverslip interface which affect emission properties [43, 47, 48]. The non-radiative processes, which compete with fluorescence emission, can introduce variations in *ϕ*, as observed in Förster resonance energy transfer and fluorescence lifetime imaging (FLIM) studies [49]. However, in the majority of cases, when the fluorophore is not intentionally employed as an environmental probe, its quantum yield can be considered constant within the sample. The brightness is directly linked to the detected fluorescence photon flux *F*_*det*_ from a single emitter, expressed as:

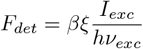

where *ξ* is the collection efficiency and *I*_*exc*_ is the laser irradiance at the molecule’s position. The collection efficiency *ξ* depends spectrally on the microscope’s optical components (objective, dichroic mirror, detection filter, detector sensitivity) but is relatively insensitive to the emitter position. It can be considered homogeneous across the field of view (FOV) for standard aplanatic microscope setups.

The ability to retrieve the molecular brightness from measurements of *F*_*det*_ requires precise knowledge of the irradiance profile *I*_*exc*_ across the field of view (FOV). In SMLM, a uniform illumination is highly desirable to ensure consistent localization precision over the entire FOV. A simple approach to achieve this is to use a wide Gaussian beam, where the irradiance can be approximated as a flat-top-like illumination. However, this method results in significant light loss, as a substantial portion of the beam must be discarded to maintain uniformity, ultimately restricting the usable FOV. Alternatively, beam shapers [50] or multimode fibers [51, 52] can be employed to generate more homogeneous excitation profiles.

In our setup (see Fig. S1), we use a dynamic illumination technique called ASTER (Adaptable Scanning for Tuneable Excitation Regions) [53], which employs rapid scanning of a small Gaussian beam to create a large and spatially uniform illumination area over the duration of the camera exposure. This technique ensures a homogeneous excitation field while remaining compatible with sectioning methods. The excitation is preferentially performed in HiLo modality [54], which effectively reduces background fluorescence while maintaining constant irradiance along the optical axis.

Beyond uniform excitation, detection homogeneity across the FOV must also be considered, particularly concerning the sample depth relative to the coverslip. Fluorophores emit supercritical angle fluorescence (SAF), which can alter flux measurements for molecules located within a distance of approximately half the emission wavelength (Fig. S1e) [47, 48]. For these fluorophores, SAF emission must be selectively removed by introducing a physical mask in a relay plane of the back focal plane of the objective (see Fig. S1). This ensures that only under-critical angle fluorescence (UAF) is collected, which remains independent of the fluorophore’s axial position [55]. For fluorophores located beyond half the emission wavelength *≈*300 nm) from the coverslip, the detected fluorescence remains axially homogeneous, allowing for direct compatibility with standard SMLM detection systems.

The collection efficiency *ξ* can be estimated by accounting for the impact of optical elements in the detection path, such as optical filters, dichroic mirrors, and the cameras spectral response, on the detected fluorescence signal. This spectral sensitivity provides an additional degree of freedom that can be optimized to enhance photon flux demixing. To quantitatively describe this effect, we define the detected brightness *β*_*det*_ = *ξβ*, which incorporates the detection efficiency of our setup (see Table 1). In the standard case of a cell labeled with two fluorophores of different brightness for multiplexing (see Fig. 1a), the goal is to distinguish these fluorophores by measuring their photon flux. However, a major challenge arises from the fact that fluorophore blinking follows a Poisson process, meaning that the duration of the ON state (*t*_ON_) varies randomly from one event to another.

**Fig. 1.**
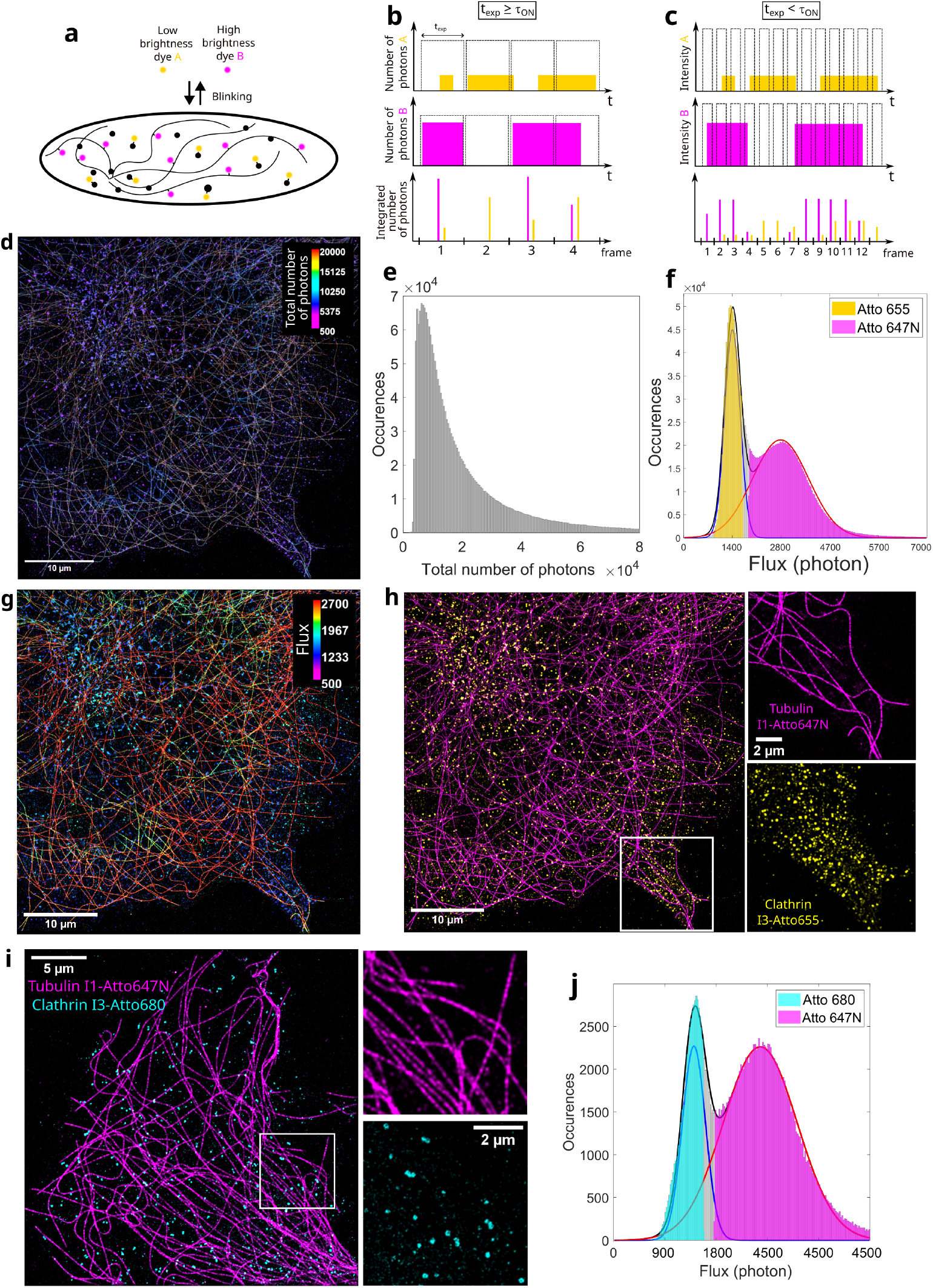
a: Cells labelled for simultaneous SMLM imaging of two targets with low and high brightness dyes. b: Chronogram illustrating molecule blinking undersampling when the exposure time *t*_*exp*_ is higher or equal to the mean ON time. c: Chronogram illustrating molecule blinking oversampling when the exposure time *t*_*exp*_ is chosen lower than the mean ON time. The flux is determined by merging localizations of the same molecule, excluding transient binding frames. d-h: COS7 cell labelled for microtubules and clathrin-coated pits with two orthogonal DNA-PAINT P1 and P3 docking strands, imaged with I1-Atto647N and I3-Atto655 respectively. d: Intensity colour-coded localization image representing the total number of photons for each event. e: Intensity histogram from Fig. 1d. f: Flux histogram following oversampled detection associated to Fig. 1g, with two gaussian-fitted peaks and a 70% specificity threshold. g: Flux colour-coded molecule image. h: Two colour-coded image after classification: microtubules (pink) and clathrin (yellow). i: COS7 cell labelled for clathrin-coated pits (cyan) and microtubules (magenta) with two orthogonal DNA-PAINT P1 and P3 strands, imaged with I1-Atto647N and I3-Atto680 respectively. j: Flux histogram associated to Fig. 1i after merging, with two gaussian-fitted peaks and a 70%4specificity threshold.

In conventional SMLM acquisition, the exposure time *t*_exp_ is typically set to match the average ON time of fluorophores (*t*_ON_). While this optimizes localization precision from a single image, it has a significant drawback: due to the stochastic nature of *t*_ON_, a fluorophore may turn ON or OFF at any moment within *t*_exp_. As a result, the number of detected photons per image does not directly reflect the fluorophore’s photon flux, since the actual emission duration may be shorter than *t*_exp_ or extend over multiple images. This leads to a large dispersion in the measured intensities, making it ineffective for distinguishing fluorophores (see Fig. 1b).

To overcome this issue, *t*_exp_ must be reduced so that *t*_exp_ ≪ *t*_ON_, as illustrated in Fig. 1c. This strategy enables oversampling of each blinking event across multiple consecutive frames. It then becomes possible to accurately identify when a fluorophore turns ON, track its fully ON state over multiple frames, and detect its extinction. By focusing only on the intermediate frames, where the fluorophore remains entirely ON, one can reliably measure its photon flux and deduce its brightness. All blinking events detected in fewer than three frames are discarded. To associate localizations from the same blinking event, we merge detections appearing within a predefined optimized radius across consecutive frames (see Methods and Fig. S2). This improves flux estimation by consolidating localizations into single blinking events, reducing measurement variability and enhancing the effective differentiation of fluorophores with different brightness.

## 3 Results

To experimentally validate brightness demixing, we used DNA-PAINT, an optimal framework for single-molecule detection due to its high and temporally stable photon yields. DNA-PAINT ensures long ON times (*t*_ON_ *≈* 150 *−* 200 ms) (see Fig.S3), allowing for flexible acquisition settings and over-sampling of blinking events. Here, we set the exposure time to *t*_exp_ = 50 ms, ensuring each blinking event is captured across multiple frames. This enables accurate flux estimation by analyzing only the frames where the fluorophore remains fully ON. For demonstration, we selected DNA-PAINT imagers Atto647N and Atto655, two commonly used far-red dyes with nearly identical spectral emissions (emission maxima separated by 16 nm) but a detected brightness ratio of 2.8 (see Table 1). To highlight the importance of flux measurement and the limitations of relying solely on total photon counts, we generated a long-exposure equivalent measurement by integrating all detected photons from the moment a fluorophore turns ON until it switches OFF. This approach mimics a single long exposure, accumulating photons over each blinking event. When localizations are color-coded according to their total photon count, microtubules and clathrin-coated pits remain indistinguishable (Fig. 1d). The corresponding histogram follows an exponential distribution, characteristic of a Poissonian process (Fig. 1e), where high variability prevents clear fluorophore discrimination. By contrast, oversampling blinking events and extracting photon flux provide a robust demixing strategy. The two fluorophore populations are now clearly separated in the flux histogram (Fig. 1f). The improved contrast is also evident in the flux color-coded image (Fig. 1g), where the two distinct biological structures emerge. Additionally, many localizations corresponding to background noise are removed, likely originating from short misbinding events or diffusing imagers in the detection focal plane.

To classify the two fluorophore populations, the flux histogram (Fig. 1f) is fitted with a sum of two Gaussian curves (see Methods). This fitting process allows the definition of a specificity threshold, enabling the exclusion of molecules with flux values falling between the two peaks. Based on this classification, microtubules and clathrin-coated pits are separated into two distinct images, as shown in Fig. 1h. To assess the accuracy of this classification, we quantify the proportion of molecules assigned to the incorrect population by calculating the cross-talk. The degree of cross-talk is dictated by the overlap between the photon flux distributions of the two fluorophores. Given that microtubules and clathrin-coated pits have distinct morphologies, we can estimate cross-talk by examining regions with filamentous microtubules, where only microtubule-associated fluorescence is expected, and vice versa for clathrin-coated pits. This estimation inherently includes both labeling-based cross-talk and misclassification due to similar photon flux values, as these two contributions cannot be disentangled. With a specificity threshold of 70% (see Methods for details), the estimated cross-talk (Fig. S4b) is 6.4% in microtubule-rich regions, corresponding to the fraction of Atto655 molecules erroneously assigned in this wrong area. In clathrin-rich regions, 4.5% of the detected Atto647N molecules are misclassified.

To assess brightness demixing (BD) in comparison to standard method, we acquired images of the same cell using spectral demixing. At a fixed specificity threshold of 70%, brightness demixing exhibited only slightly higher cross-talk than spectral demixing (6.4% vs. 4.5% for microtubules and 4.5% vs. 3.6% for clathrin) (Fig. S4d). However, spectral demixing discarded a significantly larger fraction of localizations (47.6% for SD compared to 20% for BD). This is because splitting photons into two detection channels in spectral demixing reduces the number of detected localizations, limiting the efficiency of localization pairing. Fig. S5a shows that many of the discarded localizations in spectral demixing are actually part of relevant structural information, which suggests that valuable data is lost in the process. In contrast, brightness demixing, by collecting all photons in a single channel, improves the likelihood of detecting even low-intensity blinking events. Fig. S5b demonstrates that the single-frame localizations discarded during the merging process in brightness demixing are often artifacts or background signals, originating from short misbinding events or diffusing undocked imagers. Brightness and spectral demixing could also be combined to further expand the number of parameters available for fluorophore discrimination, potentially enabling the simultaneous imaging of a larger set of dyes.

Brightness demixing is not limited to a specific pair of fluorophores but is a generalizable approach that can be applied to various dye combinations. To illustrate this, we demonstrate its applicability with an alternative fluorophore pair. By replacing Atto655 with Atto680, we increased the detected brightness ratio to 4.25. Fig. 1i shows microtubules and clathrin-coated pits imaged in a COS7 cell with Atto647N and Atto680, respectively. Brightness demixing reveals two distinct peaks in Fig. 1j and achieves comparable cross-talk values of 2.2% and 6.2% for microtubules and clathrin, respectively (Fig. S6a). We further verified that the flux histograms of these dyes (Atto647N, Atto655, Atto680) remained consistent across different targets, such as microtubules and clathrin, with only slight variations (Fig. S7). This confirms that the photon flux is independent of the target.

We take a step further by demonstrating the simultaneous demixing of three labeled targets using brightness demixing. In Fig. 2a, microtubules, clathrin-coated pits, and vimentin filaments are simultaneously imaged with Atto647N, Atto655, and Atto680, respectively. Despite a detected brightness ratio of only 1.5 between Atto655 and Atto680, Fig. 2b shows that three peaks can still be distinguished. To minimize cross-assignment errors, careful selection of demixing thresholds is required. This leads to a higher rejection rate of unspecified moleculesaround 30% in each demixed channel (Fig. S6b). This choice maintains a low cross-talk level of around 10% for each population (Fig. S6b), Alternative thresholding strategies would be beneficial, and this is a common issue with spectral-based approaches as well [34], when the number of populations to be disentangled increases. Brightness demixing is not inherently limited to three fluorophores and can be extended further by optimizing brightness contrast.

**Fig. 2.**
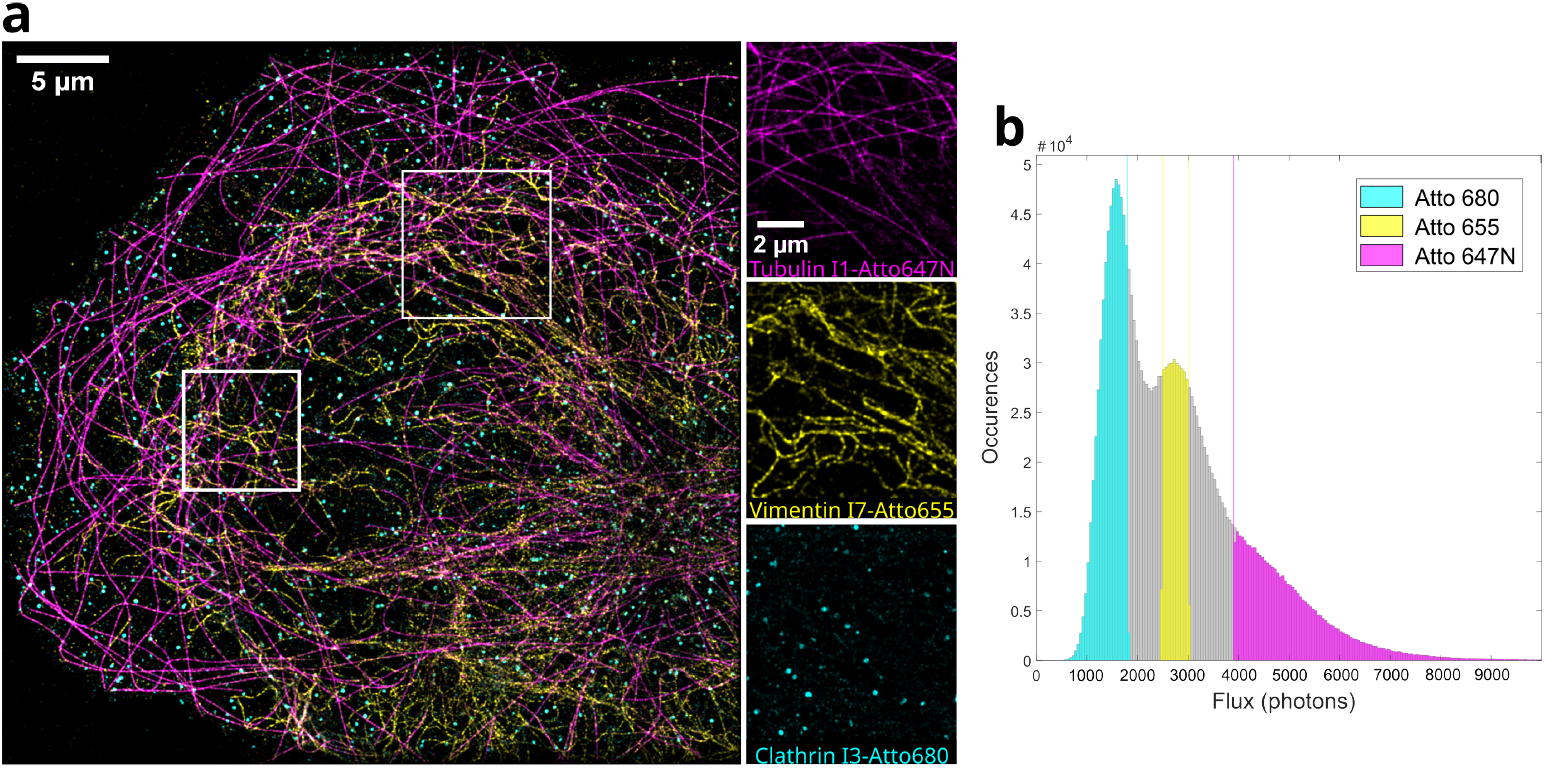
a: COS7 cell labelled for microtubules (magenta), clathrin (cyan), and vimentin (yellow) with P1, P3, and P7 strands, imaged with I1-Atto647N, I3-Atto680, and I7-Atto655 respectively. b: Flux histogram from Fig.2a after merging, with two gaussian-fitted peaks and arbitrary bounds.

We now explore the integration of brightness demixing into 3D SMLM imaging strategies, ensuring compatibility with existing methodologies. We first implemented an astigmatism-based approach, which requires only the insertion of a cylindrical lens into the detection path (Fig. 3a) [56, 57]. A key challenge is accounting for the enlarged PSF, which can affect flux estimation during photon counting (see Fig. S8). Fig. 3b displays brightness-demixed images of microtubules and clathrin-coated pits, with their respective axial localization. This approach achieves cross-talk levels comparable to those in 2D SMLM, as measured in Fig. S6c with 6.6% in the microtubule-rich area and 4.8% in the clathrin-coated pit region at a specificity threshold of 75%.

**Fig. 3.**
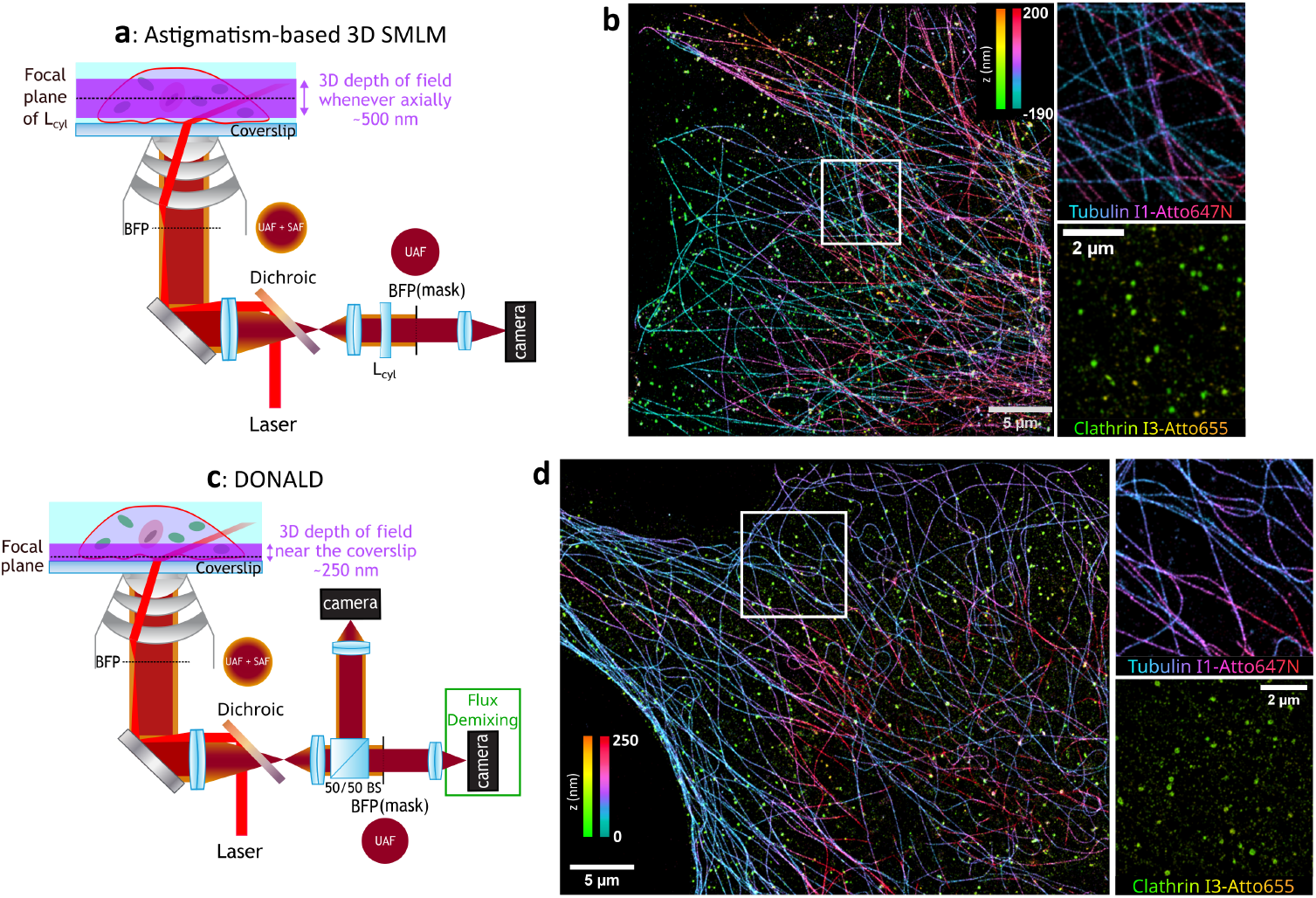
a: Setup for astigmatism-based 3D SMLM with a cylindrical lens. c: Setup for SAF-3D SMLM with DONALD configuration. b-d: COS7 cell labelled for microtubules and clathrin coated pits with P1 and P3 strands, imaged with I1-Atto647 and I3-Atto655 respectively. Both orthogonal color bars indicate (b) the relative axial position with respect to the cylindrical lens focus and (d) the absolute axial position relative to the coverslip.

Supercritical angle fluorescence (SAF) emission provides an alternative 3D imaging strategy that is inherently compatible with brightness demixing [43, 44]. In our DONALD configuration [43] (Fig. 3c), axial positioning and photon flux measurements are extracted simultaneously from the same acquisition. This is achieved by introducing a 50:50 beamsplitter, which separates the fluorescence signal into two channels. The first channel, referred to as the EPI channel, remains unaltered and contains both undercritical (UAF) and supercritical (SAF) emission components. A physical mask placed in a relay plane of the back focal plane selectively removes SAF emission, ensuring that the second channel contains only UAF. This setup allows brightness demixing to be applied directly within the UAF channel, without modifying the optical configuration. The axial position of molecules is determined by comparing the photon counts in the two channels for each localization event (Fig. S5f). Compared to astigmatism-based 3D techniques, DONALD localization offers absolute axial positioning free from sample drift, requires no calibration, and maintains a consistent PSF shape for more reliable photon flux estimation. Fig. 3d shows the brightness-demixed image of microtubules and clathrin-coated pits obtained from the UAF channel, along with their absolute axial positioning. At a specificity threshold of 80%, the cross-talk is 7.6% in the microtubule area and 8.4% in the clathrin area (Fig. S6f), comparable to 2D experiments. These results demonstrate that brightness demixing can be seamlessly integrated into existing 3D SMLM techniques, preserving their benefits while providing an additional multiplexing capability without requiring complex modifications to the optical setup.

## 4 Discussion

We have demonstrated that brightness can be used as a new property for robust simultaneous identification of multiple dyes in SMLM. Brightness demixing requires minimal modifications to existing SMLM microscopes, making it particularly well-suited for integration into compact and streamlined platforms. Only a uniform illumination and a single detection channel are necessary, with an adapted acquisition rate to extract the single-molecule flux signature. This one-channel setup avoids photon loss typically associated with splitting the detection light, as required in spectral demixing methods [31, 34] and spectrally-resolved techniques [35–37].

Our current implementation of brightness demixing is based on a single-wavelength excitation, enabling simultaneous imaging of multiple targets with a brightness contrast down to 1.5. This allows for a reduced acquisition time while ensuring compatibility with spectrally overlapping fluorophores, thereby minimizing chromatic aberration. Compared to spectral demixing, brightness demixing preserves a higher number of localizations and avoids constraints associated with channel splitting. We also demonstrated straightforward compatibility with 3D SMLM techniques such as DONALD and astigmatism-based microscopy, maintaining cross-talk levels comparable to 2D imaging. In particular, supercritical angle fluorescence (SAF)-based 3D SMLM methods [43, 44] inherently require a two-camera setup to retrieve axial information. Introducing spectrally resolved detection in such systems would lead to a four-channel configuration, complicating optical alignment and increasing data processing challenges. In contrast, brightness demixing preserves the simplicity of the imaging setup while enabling multi-target imaging. Additionally, by utilizing all detected photons in a single channel, this approach can accommodate larger fields of view (*>* 100*µm*^2^), facilitating simultaneous imaging of multiple cells, as demonstrated in Fig.S9. Compared to alternative single-camera strategies using composite PSFs for multicolor imaging [9, 10, 16], brightness demixing operates with the same blinking density as conventional 2D SMLM experiments. Furthermore, its reliance on a single laser excitation minimizes chromatic aberration, making it an attractive alternative for high-throughput multi-target imaging.

DNA-PAINT is rapidly evolving, particularly to address initial limitations related to high background from unbound imagers, as well as constraints in imaging speed and throughput [27, 28, 58–61]. Reduced acquisition times are now achievable using commercial FAST-PAINT imagers, which feature shorter binding times and reduced background noise. We demonstrate in Fig.S10 that brightness demixing remains compatible with these newer imagers, using a frame rate of 10 ms and a total acquisition of 80,000 frames in 13 minutes. Current developments in brightness demixing still depend on the availability of dyes optimized for DNA-PAINT, which were initially designed for spectral separation rather than brightness contrast [62]. However, by demonstrating that brightness provides an additional contrast mechanism in SMLM, we anticipate that new dye development efforts will focus on fluorophores with identical spectral properties but distinct brightness, further facilitating their classification [63, 64]. Brightness demixing, by enabling at least three simultaneous targets in a setup traditionally limited to sequential acquisition, such as Exchange-PAINT [24, 27, 28], will significantly accelerate and enhance multi-target imaging for cellular spatial proteomics.

Beyond its role in multiplexing through brightness demixing, precise photon flux measurement opens new avenues for probing quantum yield variations, which are inherently linked to fluorescence lifetime. This principle is central to fluorescence lifetime imaging microscopy (FLIM), providing insights into local molecular interactions and environmental changes. However, applying such an approach in SMLM requires the development of novel dyes with tailored blinking properties to enable accurate flux quantification. Future efforts in engineering fluorophores with controlled blinking and tunable quantum yield could bridge brightness demixing with functional imaging, further expanding molecular-scale investigations. Moreover, in the context of live-cell super-resolution microscopy [65– 68], where dynamic and local chemical interactions are crucial, these measurements could provide valuable tools for studying biological mechanisms in real time.

## 5 Methods

### 5.1 Biological samples preparation

#### Cell Culture

COS-7 and HeLa cells were grown in DMEM with 10% FBS, 1% L-glutamin and 1% penicillin/streptomycin (Life Technologies) at 37^*°*^C and 5% CO2 in a cell culture incubator. For the U2OS cell culture, DMEM was replaced by McCoys 5A medium. After two days in incubator, they were seeded at medium confluence on cleaned and round 25 mm diameter high resolution 1.5” glass coverslips (Marienfield, VWR).

#### Cell fixation for microtubules, clathrin and vimentin imaging

After 24 hours, the cells were washed one time with sterile PBS 1X (Invitrogen, 003000). An extraction solution (0.25% Triton, 0.1% glutaraldehyde in PEM) is added for 45s then a fixation solution (0.5% glutaraldehyde, 0.25% Triton in PEM) for 10 min at 37^*°*^C followed by a reduction solution (0.1mM NaBH4) for 7 min. The PEM solution concentration is 80 mM PIPES, 5 mM EGTA, and 2 mM MgCl2 adjusted to pH 6.8 with KOH 6M. Extraction and fixation solutions were pre-warmed at 37^*°*^C for 15 minutes at least. Cells were then washed 3 times in PBS 1x.

#### Cell labeling for DNA-PAINT imaging

For the saturation step, cells are blocked for 30 min in PBS + 3% BSA. Primary antibodies are introduced simultaneously for multi-target imaging. The cells were incubated 1h at 37^*°*^C with primary antibody (see Table 3) in PBS + 3% BSA. Secondary antibodies are introduced simultaneously for multi-target imaging. This was followed by three washing steps in PBS+3% BSA and incubation for 1h at 37^*°*^C with secondary antibodies combined with DNA-PAINT strand (see Table 4) diluted in antibody buffer (Massive Photonics). Three more washes with PBS 1x are performed. Post-fixation is performed with firstly formaldehyde 3.7% in PBS, then three washes with PBS 1x, and secondly NH4Cl (2.7 g/L) in PBS followed by three washes with PBS 1X. The coverslips are stored in washing buffer (Massive Photonics).

### 5.2 SMLM microscope

The sample was imaged through a 100x 1.49 NA APO TIRF WD 0.13-0.20 oil objective lens on a Nikon Ti2 Eclipse (Nikon France SAS, Champigny sur Marne, France) inverted microscope associated to a SAFe 360 detection module (Abbelight, Cachan) combined to two sCMOS Flash 4 v3 camera (Hamamatsu, Japan). Uniform and large field of view excitation is provided through an ASTER module [53], which included 639 nm (ERROL laser 500 mW) lasers and a quad band dichroic/emission filter (Semrock 405/488/532/640 refs. FF01-446/510/581/703-25 and Di03-R405/488/532/635-t1-25×36). The acquisition is driven by Neo software (Abbeligth, Cachan). A physical mask is placed in a relay plane of the back focal plane of the objective to block the SAF emission. The alignment of the mask is made by adding a Bertrand lens. For Brightness demixing, only one camera is used.

For astigmatism-based 3D SMLM, the point spread functions were shaped by inserting a cylindrical lens with a focal length of -25m in the optical pathway between the dichroic mirror and the camera. Three-dimensional astigmatism-based acquisitions were fitted (Gaussian fitting - MLE) for z position according to the eccentricity of the PSF using a calibration obtained on 40 nm beads (660 excitation/680 emission wavelength). We account for PSF anisotropy during the photon counting step, using an integration mask over 17 pixels (see Fig.S8).

For 3D imaging based on supercritical fluorescence emission, the SAFe detection module was configured for DONALD microscopy [43], i.e. the fluorescence image is divided into two imaging paths and imaged on two cameras by a 50 − 50% beamsplitter, and intrinsic supercritical angle fluorescence (SAF) is used to extract the absolute axial position of single-molecule events. 3D DNA-PAINT analysis including drift correction, axial information calculation (SAF and astigmatism) and ratiometric measurements for spectral demixing were performed with Neo analysis software (Abbelight Cachan).

For comparison with spectral demixing acquisition, the fluorescence image is divided into two complementary spectral images by a dichroic with a cut-off wavelength at 700 nm (Chroma T700lpxr-3).

### 5.3 DNA-PAINT acquisition

For DNA-PAINT imaging, the imager strands are diluted in imaging buffer (from Massive Photonics) achieving a concentration of 0.2 to 1 nmol in the sample chamber (see Table 5). For multi-target imaging, all the imagers were added simultaneously into the imaging buffer. The illumination field is about 30 x 30 *µ*m^2^ with an irradiance of approximately 10 kW.cm^*−*2^, which corresponds to a 100 mW laser beam scanned on slighlty largera area of 35×35 *µ*m^2^ with ASTER. The excitation is in Highly Inclined and Laminated Optical sheet (HILO) [54] configuration for removing background fluorescence from unbound imagers. Exposure time is fixed at 50 ms and from 40 000 to 80 000 frames have been taken with a focus maintaining system (PFS, Nikon). For acquisition with FAST PAINT sequences, the exposure time is decreased to 10 ms.

### 5.4 SMLM data processing

For the localization data processing, the Abbelight Neo Analysis software (Abbelight, Cachan) was used to retrieve all the molecules localized per frame with their corresponding intensity. Detection of intensity peaks was done using a wavelet algorithm [69] after local means background estimation and removal with a kernel size of 15 pixels. Intensity peaks of at least twice the background with a size of 3×3 to 7×7 pixels area were considered a single-molecule blinking event and further processed for fitting using gaussian fitting with Maximum Likelihood Estimation (MLE). The photon counting step was estimated, after local means background estimation and removal with a kernel size of 15 pixels, by integration over a 9 pixels round area (1 pixel is equal to 97 nm on the sample). The influence of the integration mask size has been studied and optimized on the cross-talk value and rejection rate (see Fig.S11). The drift is corrected with a RCC algorithm (1000 images per stack). For 3D DONALD measurement, during data processing, an affine transform is applied to correct any shift between the two channels. Each localization appearing at the same position on the two cameras is assigned together with a tolerance of 700 nm on identical frames. Then the SAF ratio is calculated for each localization by using the formula: *r*_*SAF*_ = (*N*_*SAF*_ *−N*_*UAF*_)*/N*_*UAF*_, with *N*_*SAF*_ and *N*_*UAF*_ respectively the number of photons in the SAF and UAF channel. A correction to the reflection/transmission factor of the beam splitter is taken into account, which is equal to 1.22 on our setup. For spectral demixing, after data processing, two images are reconstructed from the table of localizations of each camera. Localizations with the same spatial localization with a tolerance of 700 nm on identical frames are paired, and then the photonic ratio is calculated by: *r*_*SD*_ = *N*_*refl*_*/*(*N*_*trans*_ + *N*_*refl*_), with *N*_*trans*_ and *N*_*refl*_ respectively the number of photons in the transmitted and reflected channel.

### 5.5 Processing for retrieving Flux value

After SMLM data processing as stated above, a table with all the localization is obtained. For each localization, the algorithm was designed to search the next frame in a radius of 100 nm for another localization. The influence of the association radius is shown in Fig.S2, we choose a large enough association radius to minimize the rejection rate while avoiding false association (Fig.S2a) and keeping the cross-talk and the unspecified molecules rate (accounting the rejected localization that are not classified) low (Fig.S2c). For three simultaneous targets, we decrease the association radius to 30 nm since the total concentration of imagers is higher and the localization density too. Then for each molecule, the number of frames on which the molecule is localized is measured (called ON time, see Fig.S3) and the average intensity discounting the first and last frame is calculated, giving the flux measurement. The code is available to process localization files in csv. The localization precision for each demixed channel is calculated by using NeNa algorithm [70] and is given in Table 2.

### 5.6 Likelihood probability fit and assignment

The flux histogram is fitted by the sum of two gaussians corresponding to the population of molecules A and B. The probability of localizations at a given flux to be A is given by :

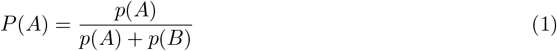

with *p*(*A*) and *p*(*B*) the proportion of molecules at this given flux, which is the value of each gaussian at this point. The crossing point of the two gaussians corresponds to a probability *P* (*A*) = *P* (*B*) = 50%. For classifying the two populations, the bounds delimiting the two populations is fixed by choosing the specificity value, which is the minimum probability of a molecule to be classified in the good population. For example, if the specificity is fixed at 60%, all localization with a probability *P* (*A*) under 0.6 will be rejected in population A. A higher specificity leads to a better classification but increases the number of rejected molecules referred to as the unspecified molecules rate (Fig.S11c). We set the specificity threshold to keep the rate of unspecified molecules low, around 10%.

### 5.7 Cross-talk analysis

Since we have selected proteins that are present on morphologically distinct organelles, this enables us to select regions wherein we have certainty of only one kind of protein being present. We assume that the proteins we are imaging don’t colocalize or at least we can find areas where the proteins are not on the same spot. We then choose to calculate the cross-talk by drawing regions corresponding to the labelled organelles. In these regions, the proportion of each population is then calculated and averaged. The number of unrecognized molecules is also calculated in these regions.

## Acknowledgments

L. Le acknowledges a PhD funding from ENS Paris Saclay, S. Sreenivas acknowledges PhD funding from UpToParis program (ESPCI, Marie Skodowska-Curie Actions). We acknowledge the use of the Centre de Photonique pour la Biologie et les Matriaux for cell culture facility within ISMO. We thank Fatiha Mahtout for support for cell cultures and fixation. We thank Abigail Illand, Lancelot Pincet and Guillaume Dupuis for helpful discussions. We thank Abbelight for the free use of NEO software. This work was supported by the AXA research fund, the ANR (grant nos. LABEX WIFI, ANR-10-LABX-24), and the European Union’s Horizon 2020 research and innovation program under grant agreement no. 871124 Laserlab-Europe for Joint Research Activity.

## Declarations

## Availability of data and materials

dataset for localization files are available on Zenodo (https://doi.org/10.5281/zenodo.14505755), complementary materials are available from corresponding author on request

## Code availability

the code for the association of events to calculate the flux and perform brightness demixing is available on Github (https://github.com/NanobioSMLM/brightnessdemixing). SMLM large raw data files (*≥*20Go) are available from the corresponding author on reasonable request.

## Authors’ contributions

L.L., E.F. and S.L.F. conceived the project, L.L. developed the practical implementations, performed the labelling, the acquisitions and analysis. S.S and S.L.F. realized preliminary experiments. L.L and E.F developed processing tools. All authors have contributed to the manuscript.

