## Supplementary Material for "Brightness demixing for simultaneous multi-target imaging in 3D single-molecule localization microscopy"

**Figure S1**: Optical setups for 2D and 3D brightness demixing.

**Figure S2**: Characterization of the DNA-PAINT imagers.

**Figure S3**: Influence of association radius during merging process.

**Figure S4**: Comparison between Spectral and Brightness demixing.

**Figure S5**: Discarded localizations map with Spectral and Brightness demixing.

**Figure S6**: Flux histogram and cross-talk estimations.

**Figure S7**: Permutation of targets with the same imager.

**Figure S8**: Influence of photon counting mask size for astigmatism-based 3D SMLM.

**Figure S9**: Brightness demixing with an extended field of view.

**Figure S10**: Brightness demixing with Fast PAINT imagers.

**Figure S11**: Influence of photon counting mask size.

**Table 1**: Calculated brightness of the different dyes used for brightness demixing.

**Table 2**: Localization precision of the different figures.

**Table 3**: Primary antibodies.

**Table 4**: Secondary antibodies.

**Table 5**: DNA-PAINT imagers.

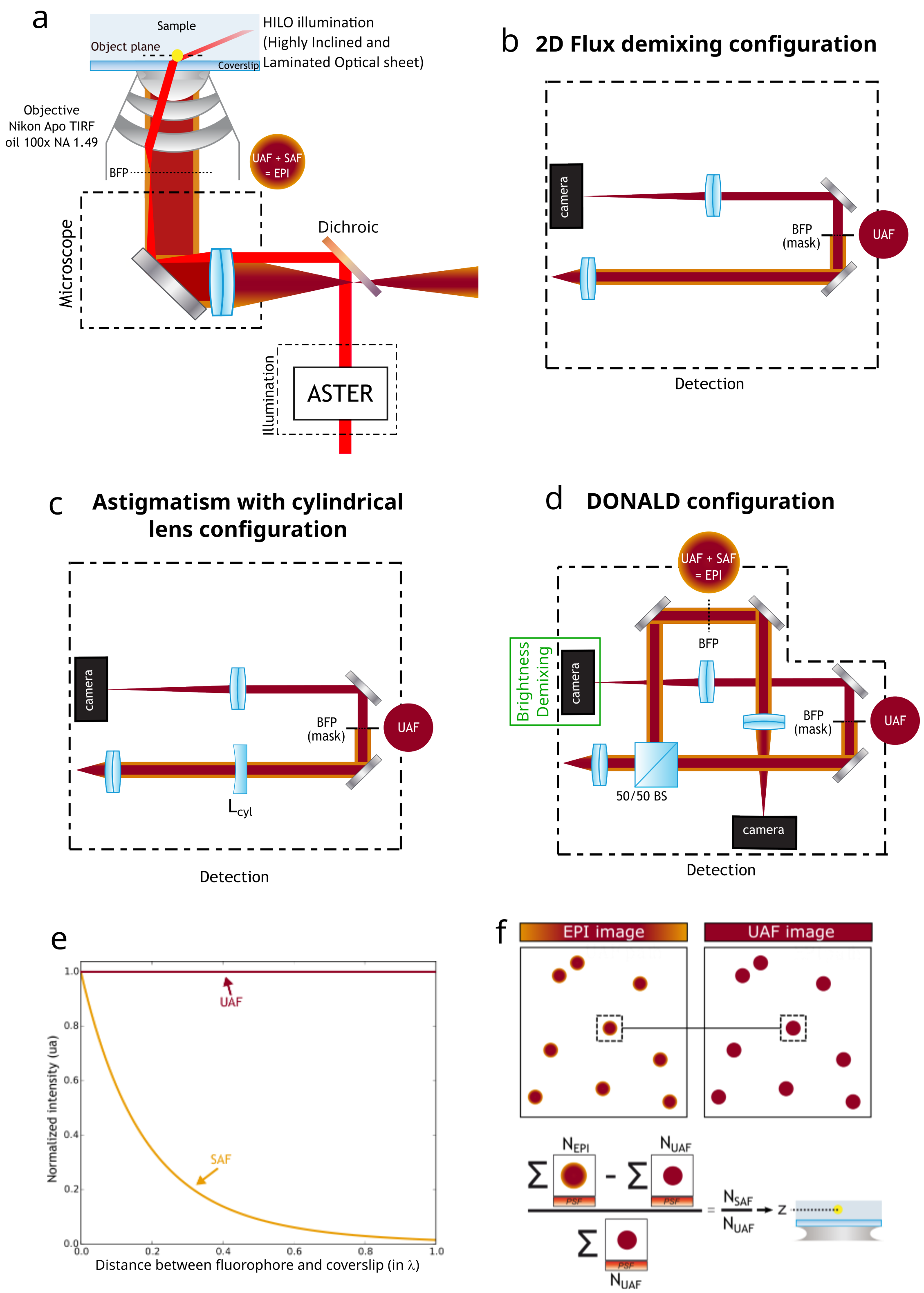


**Fig. S1** a: Optical setup of the Abbelight Safe 360 microscope. The ASTER illumination (Abbelight, Cachan) provides a homogeneous wide-field excitation by scanning a gaussian spot at the camera frame rate. The size of the excitation field can also be tuned. A motorized stage enables to change the illumination angle to a HILO configuration. b: Detection setup for single channel detection with a physical mask in the back focal plane (BFP). c: Detection setup for 3D astigmatism-based microscopy. $L_{cyl}$: Cylindrical lens with a focal length of -25 m. d: Detection setup for DONALD configuration. e: Variation of fluorescence emission as a function of the distance between the fluorophore and the coverslip. UAF (undercritical angle fluorescence) remains constant and can be used for brightness demixing, whereas SAF (supercritical angle fluorescence) varies with axial distance and enables 3D SMLM. f: Principle of DONALD microscopy, PSFs are paired from the UAF and EPI (UAF + SAF) channels to compute the SAF ratio, which is then converted into an axial distance from the coverslip.

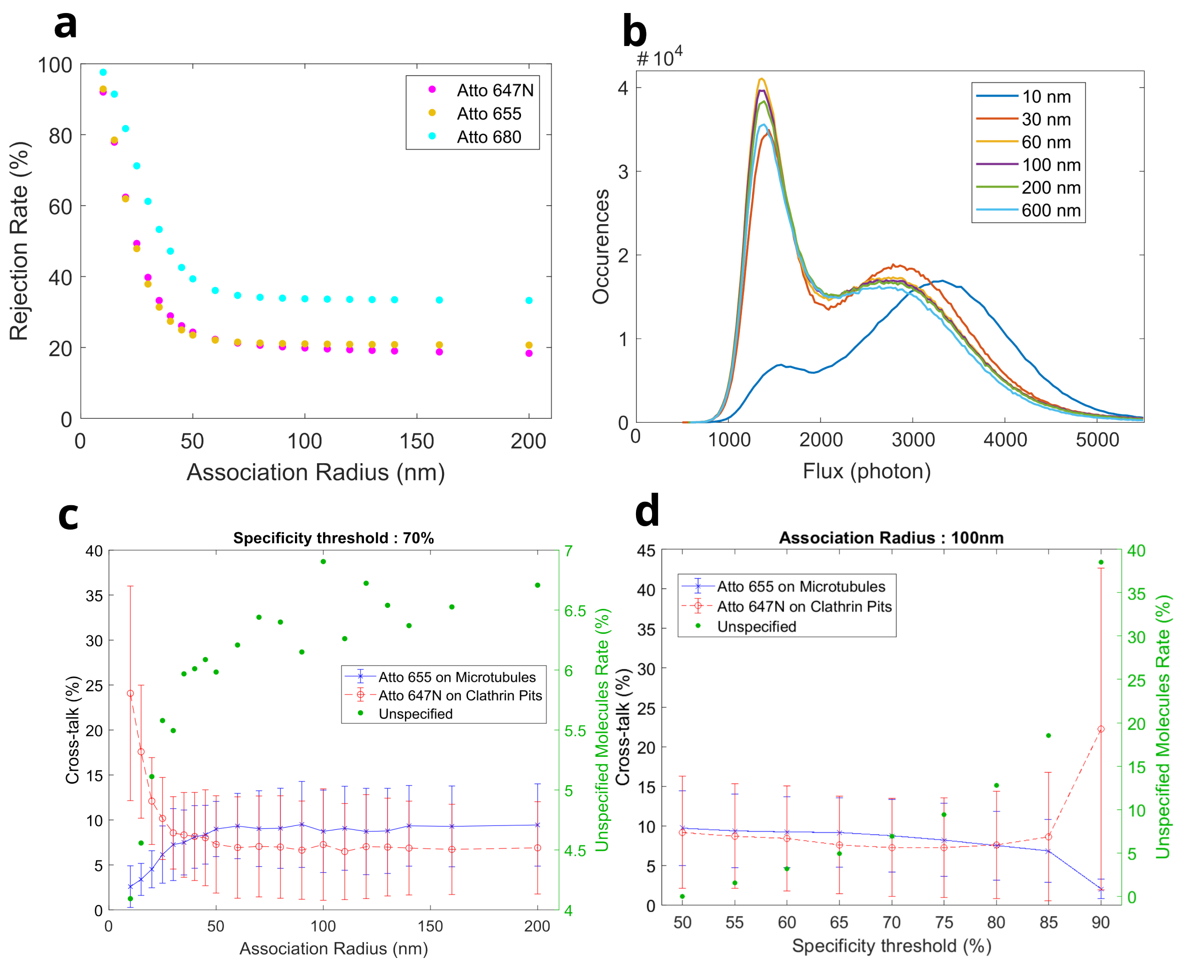


**Fig. S2** a: Rejection rate calculated during the merging process against the association radius for the three imagers Atto647N, 655 and 680. The rejection rate is the proportion of localizations found on only one or two frames and then not considered for the brightness demixing step. We can notice that the rejection rate is barely varying for an association radius greater than 70 nm. It means that the association radius is big enough for finding localizations on several continuous frames and that the localization density is low enough to avoid false association. b-d: With the dataset of Fig.1d-h. b: Flux histogram for different association radii. We can see that the association radius may affect the flux histogram (10 nm) but is stable for a bigger association radius. c: Cross-talk estimation and unspecified molecules rate against the association radius for a specificity threshold of 70%. As we can see in Figure S6b, the flux histogram for a small association radius is totally different and surely not relevant, leading then to high cross-talk for Atto647N on clathrin pits. However, for an association radius higher than 30 nm, the cross-talk is stable. d: Cross-talk estimation and unspecified molecules rate against the specificity threshold for an association radius of 100 nm. As the specificity threshold increases, the cross-talk decreases while the unspecified molecules rate increases. This is understandable as we are more specific but oddly enough, the cross-talk of Atto647N on clathrin pits increases for specificity threshold higher than 80%. Indeed, because the peak associated with Atto647N dye is wider than the one associated with Atto655, the proportion of Atto655 well demixed on clathrin pits area becomes very low at high specificity compared to Atto647N population.

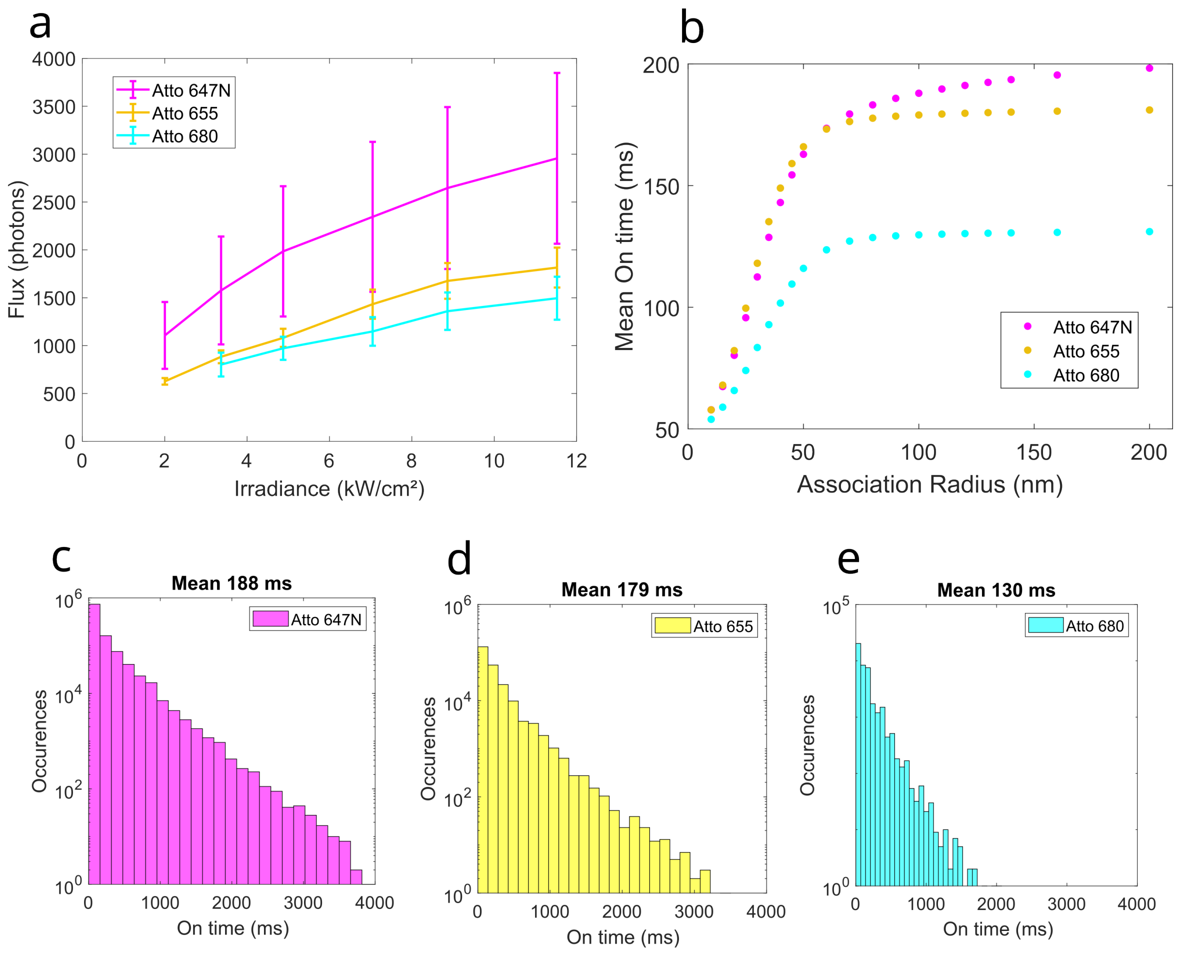


**Fig. S3** a: Flux against excitation irradiance in HILO configuration for imagers Atto647N, 655 and 680. We choose to fix the irradiance near 11 kW.cm$^{-2}$ corresponding to an average power of 100 mW in the back focal plane of the objective. b: Mean On time against association radius chosen for the merging process of localizations. The exposure time of the camera is 50 ms. We can notice that for an association radius greater than 80 nm, the mean On time barely varies, meaning that the association radius is big enough for finding localizations on several continuous frames and that the localization density is low enough to avoid false association. c, d, e: On-time histogram in log scale of imagers Atto647N, 655 and 680 respectively. The decay scales linearly, meaning that we can see that the On time distribution is mono-exponential decay.

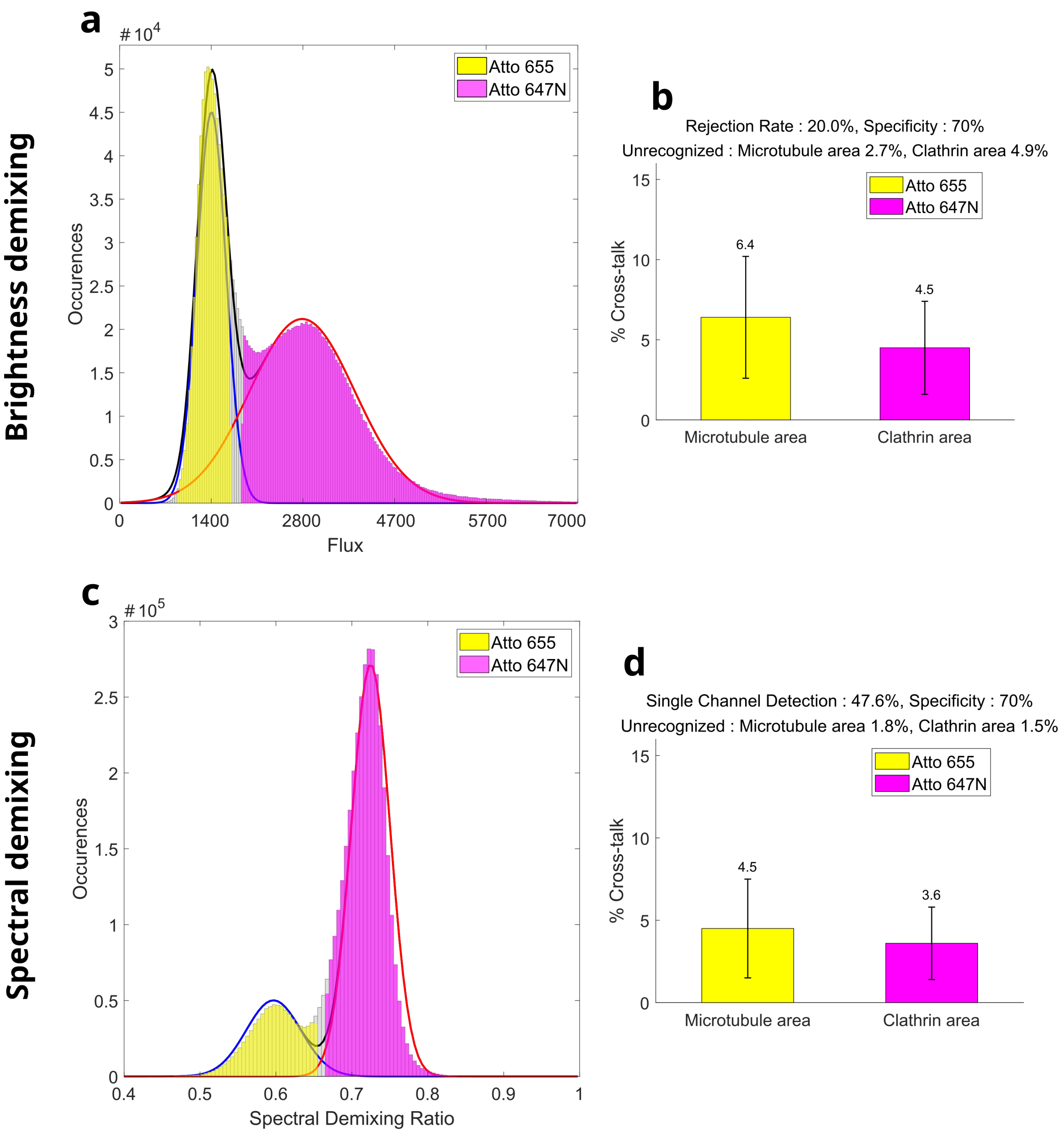


**Fig. S4** Comparison between Brightness Demixing and Spectral Demixing. COS7 cell labelled for microtubules and clathrin-coated pits with P1 and P3 docking strands simultaneously imaged with I1-Atto647N and I3-Atto655 respectively. a: Histogram of the flux distribution associated with Fig.1g. b: Cross-talk estimation for brightness demixing. c-d: Spectral demixing of the same cell in Fig.1g. c: Histogram of spectral demixing ratio. d: Cross-talk estimation for spectral demixing.

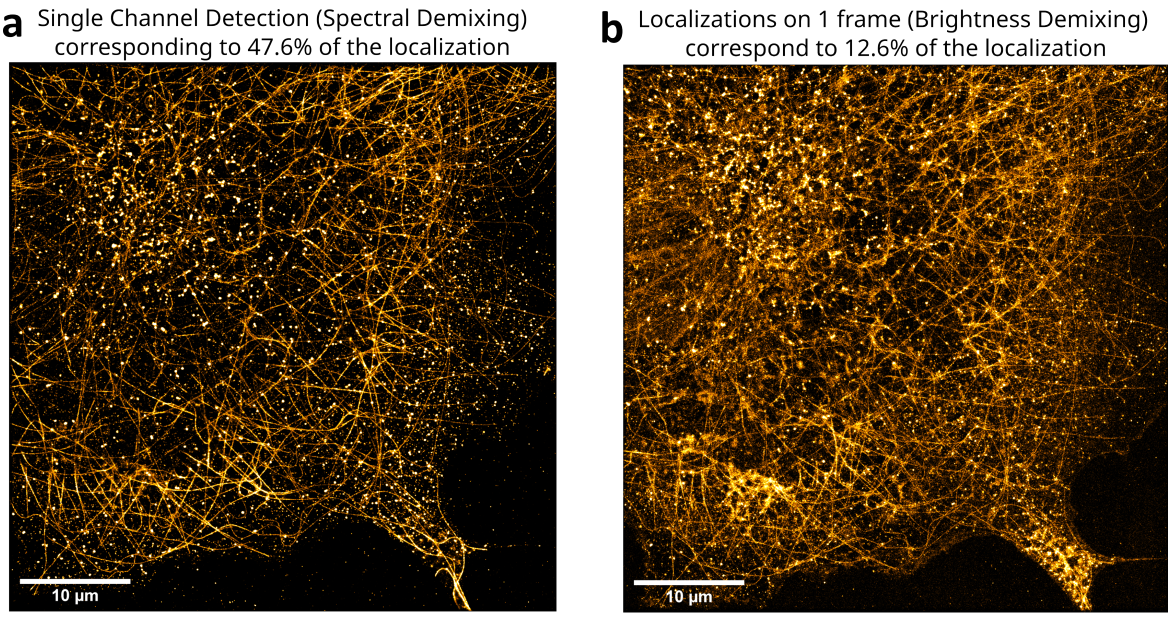


**Fig. S5** Discarded localizations during (a) Spectral Demixing and (b) Brightness Demixing. a: Localizations appearing on only one channel of the Spectral Demixing experiment from Fig.S4c, b: Localizations appearing on only 1 frame of the Brightness Demixing experiment from Fig.1d-h.

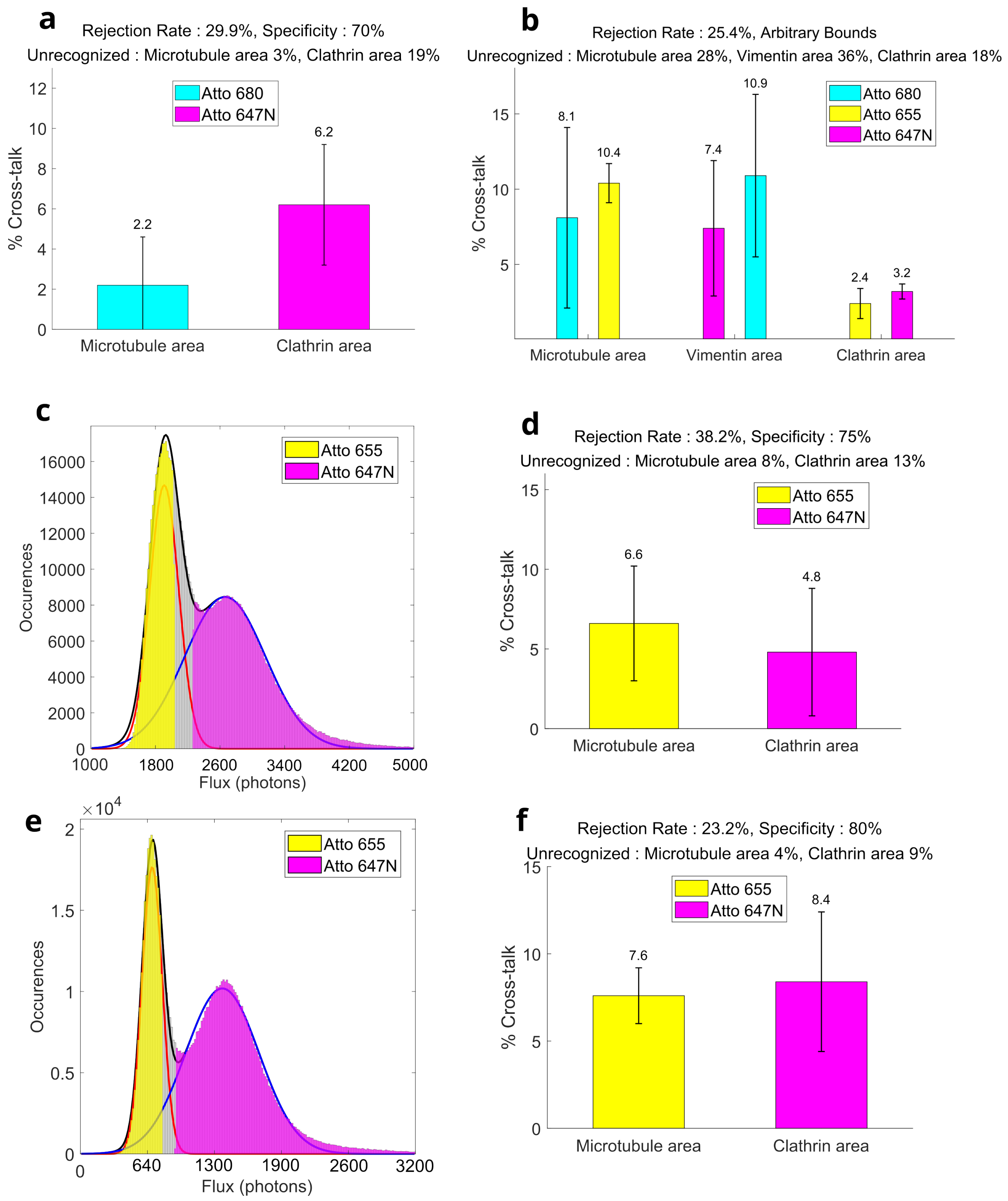


**Fig. S6** a: Cross-talk estimation associated with fig.1i. b: Cross-talk estimation associated with fig.2. c,d: Flux histogram and cross-talk estimation associated Fig.3b. e,f: Flux histogram and cross-talk estimation associated Fig.3d.

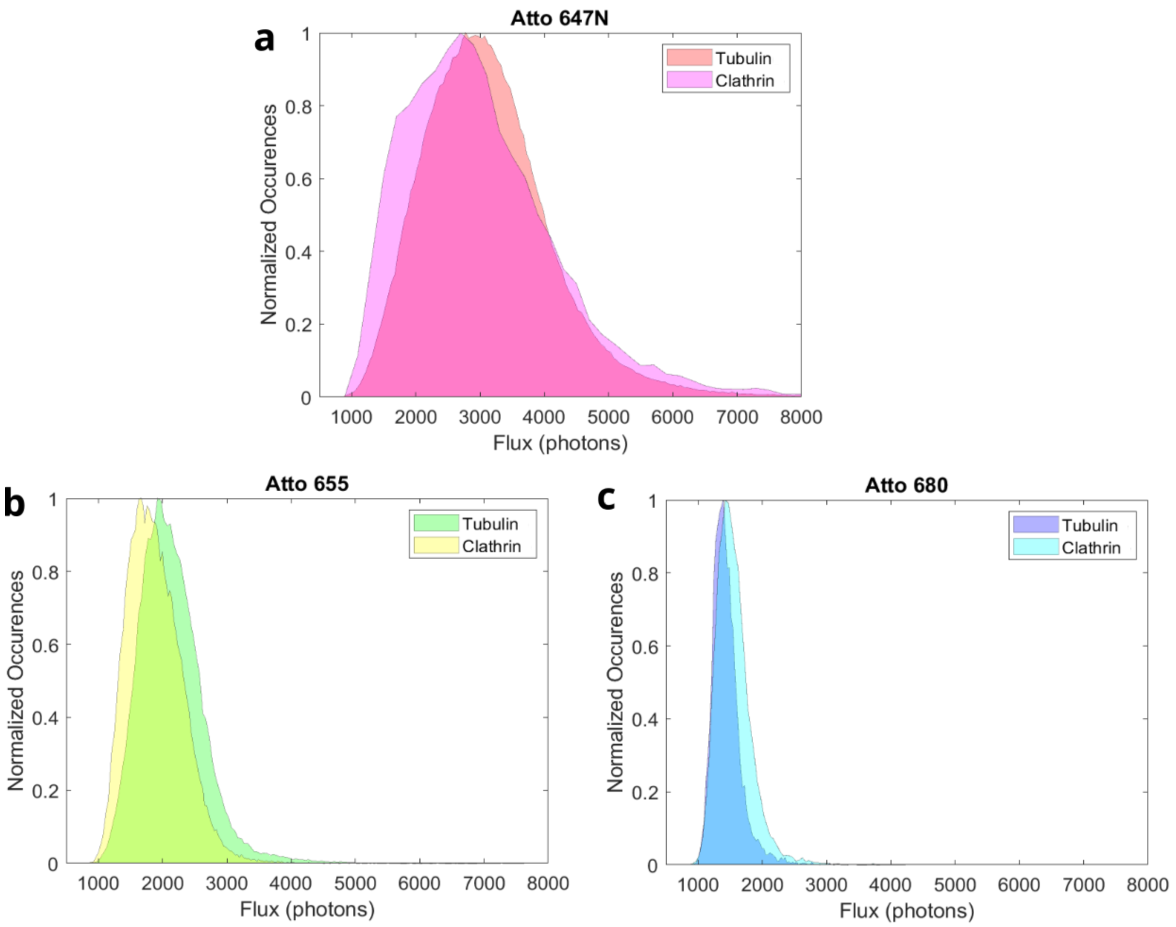


**Fig. S7** Flux histograms of Atto 647N, 655, and 680 imaging $\alpha$-tubulin or clathrin coated pits. The I1-Atto 647N, I3-Atto 655, and I3-Atto 680 imagers were used for DNA-PAINT imaging. For each imager, the docking strand remained the same when switching the target to be imaged, but different antibodies were used.

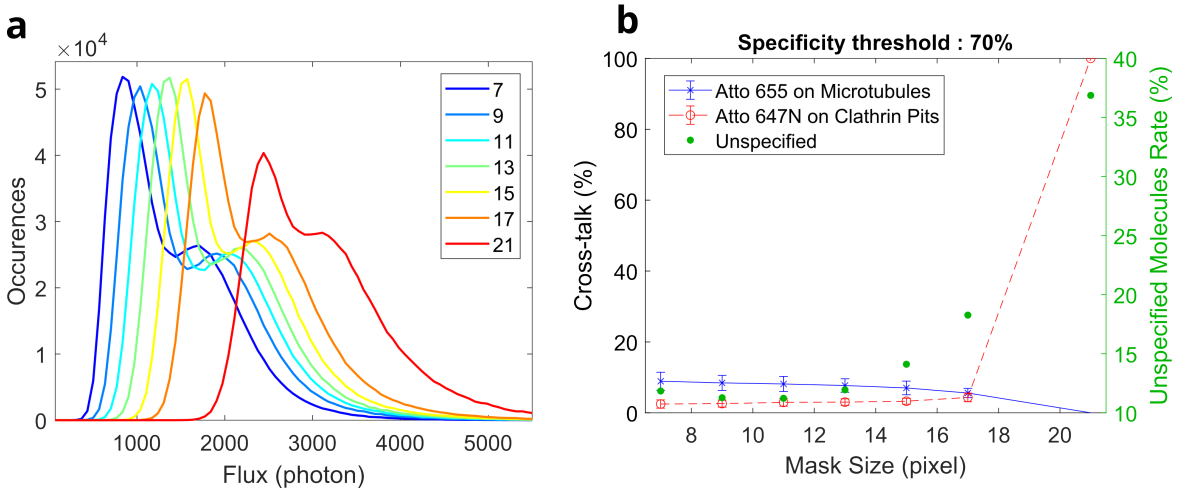


**Fig. S8** Influence of photon counting mask size on cross-talk and rejection rate for astigmatism-based 3D SMLM. a: Flux histogram profile for different photon counting mask sizes with dataset of Fig.3b in the case of astigmatism-based 3D SMLM with a cylindrical lens. b: Evolution of cross-talk as a function of photon counting mask size. The rate of non-specific molecules is shown in green, representing the proportion of molecules rejected because their probability of belonging to one of the populations is below the specificity threshold.

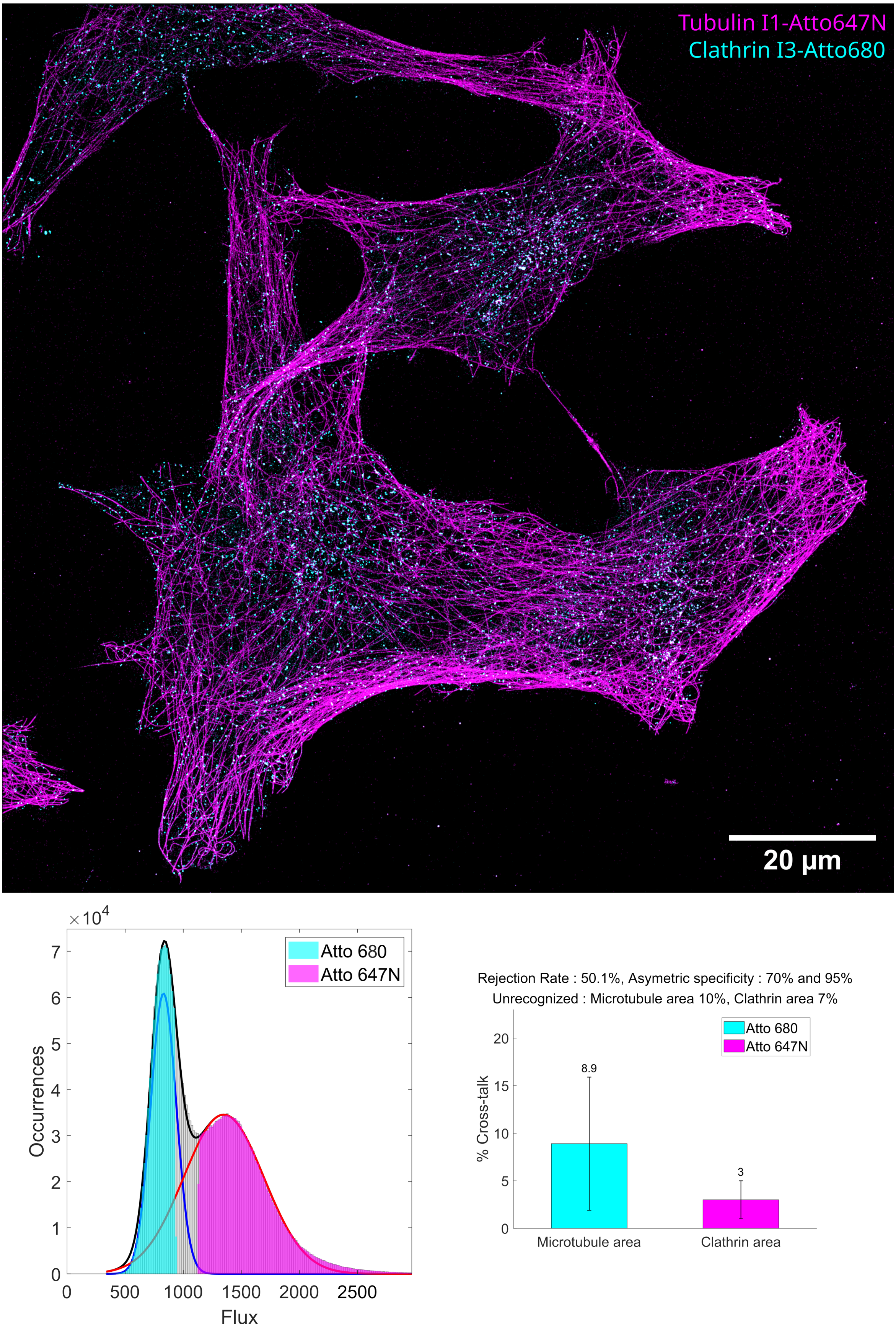


**Fig. S9** Brightness demixing with an extended field of view of more than 100 $\mu m$. COS7 cell labelled for microtubules and clathrin coated pits with P1 and P3 strands, imaged with I1-Atto647N and I3-Atto680 respectively. The flux histogram is represented with the cross-talk estimation. This figure demonstrates that brightness demixing can be performed on a wide field of view over 100 microns to image several cells. Irradiance: 5 kW.cm$^{-2}$

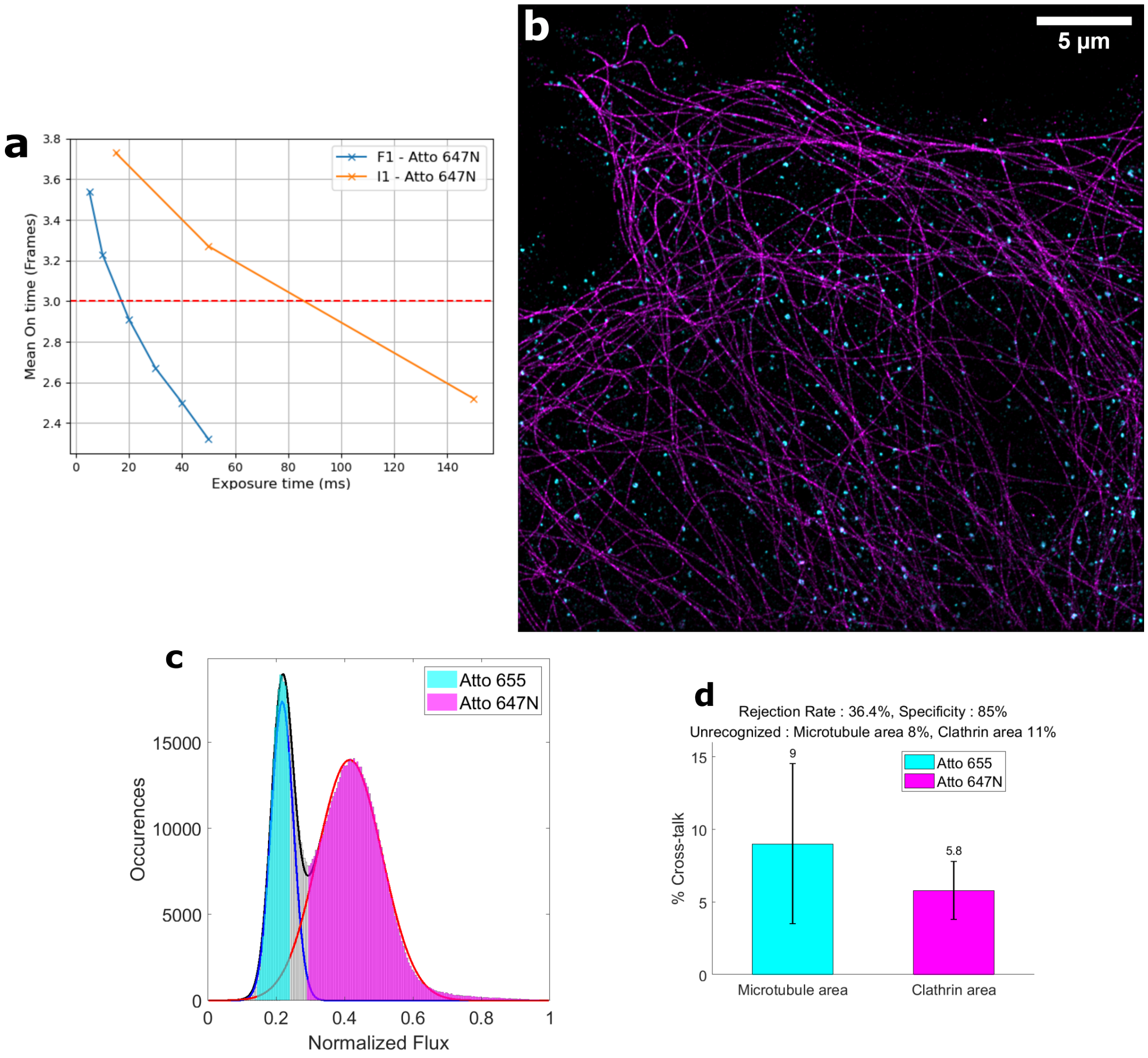


**Fig. S10** Brightness demixing with Fast PAINT imagers. a: Mean On time against the exposure time for regular PAINT (I1) and Fast PAINT (F1). The red dashed line marks the threshold at 3 frames, representing the minimum required to oversample the blinking event. Since Fast PAINT exhibits faster blinking dynamics, the exposure time was consequently reduced to 10 ms. b: COS7 cell labelled for microtubules and clathrin coated pits with F1 and F2 strands, imaged with F1-Atto647N and F2-Atto655 respectively. c-d: The flux histogram is represented with the cross-talk estimation.

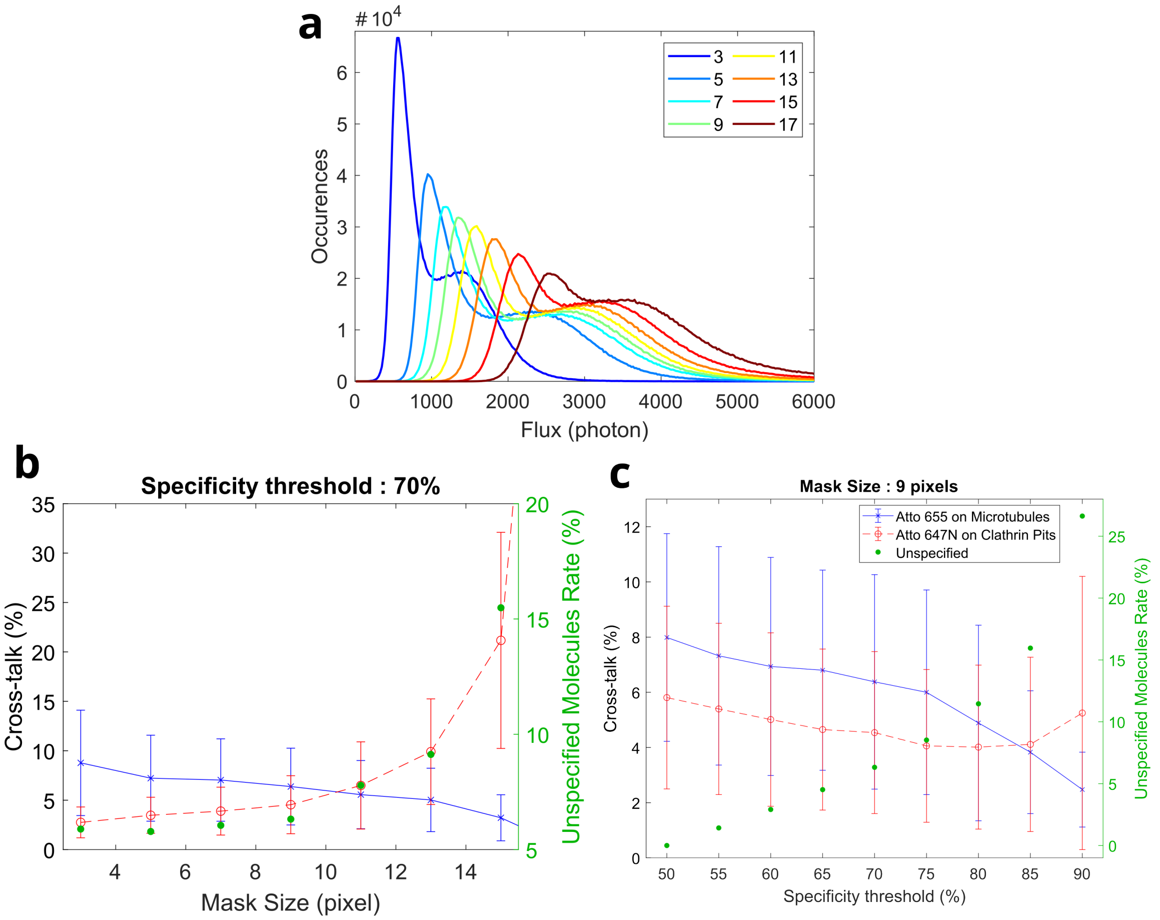


**Fig. S11** Influence of photon counting mask size on cross-talk and rejection rate. a: Flux histogram profile for different photon counting mask sizes with dataset of Fig.1d-h. b: Evolution of cross-talk as a function of photon counting mask size. The rate of non-specific molecules is shown in green, representing the proportion of molecules rejected because their probability of belonging to one of the populations is below the specificity threshold. c: Evolution of cross-talk against the specificity threshold.

| Dye | Brightness $\beta$ $(.10^4 M^{-1}.cm^{-1})$ | Detection Efficiency $\xi$ | $\beta_{det} = \xi.\beta$ $(.10^4 M^{-1}.cm^{-1})$ |
|---|---|---|---|
| Atto 647N | 8.7 | 32% | 3.4 |
| Atto 655 | 2.5 | 39% | 1.2 |
| Atto 680 | 1.7 | 46% | 0.8 |

**Table 1** The detection efficiency $\xi$ is taking into account the camera detection efficiency, the objective transmission and the transmission spectra of the dichroic and detection filter. $\beta_{det}$ is the detected brightness the product of the brightness $\beta$ and the detection efficiency $\xi$.

| Data | Low Flux | | Middle Flux | | High Flux | |
|---|---|---|---|---|---|---|
| Localization precision (NeNa) | $\sigma_{xy}$ | $\sigma_z$ | $\sigma_{xy}$ | $\sigma_z$ | $\sigma_{xy}$ | $\sigma_z$ |
| 2D Atto647N/655 (Fig.1 d-h) | 7.7 | - | - | - | 7.5 | - |
| 2D Atto647N/655 (SD) (Fig.S4 c-d) | 13.2 | - | - | - | 11.4 | - |
| 2D Atto647N/680 (Fig.1 i-j) | 8.8 | - | - | - | 10.3 | - |
| 2D Atto647N/655/680 (Fig.2) | 6.9 | - | 6.7 | - | 8.3 | - |
| 3D Astigmatism Atto647N/655 (Fig.3 b) | 11.6 | 16.2 | - | - | 12.4 | 16.3 |
| 3D DONALD Atto647N/655 (Fig.3 d) | 8.8 | 22.2 | - | - | 8.8 | 22.3 |

**Table 2** Lateral and axial localization precision $\sigma_{xy}$ and $\sigma_z$ in nanometres of the different figures.

| Structure | Target | Species | IgG type | Clone | Supplier | Cat# | Dilution |
|---|---|---|---|---|---|---|---|
| microtubules | $\alpha$-tubulin | Mo monoclonal | IgG1 | B512 | Sigma | T5168 | 1:500 |
| microtubules | $\alpha$-tubulin | Mo monoclonal | IgG1 | DM1A | Sigma | T6199 | 1:500 |
| clathrin-coated pits | clathrin heavy chain | Rb polyclonal | - | - | abcam | ab21679 | 1:500 |
| intermediate filaments | vimentin | Ch polyclonal | igY | - | Encor Biotechnology | - | 1:500 |

**Table 3** Primary antibodies used for immunostaining. Mo = Mouse, Rb = Rabbit, Ch = Chicken.

| Nanobody | Target species | Name | Sequence 5'-3' | Supplier | Dilution |
|---|---|---|---|---|---|
| sdAB | mouse | P1 sequence | TTATACATCTA | Massive Photonics | 1:500 |
| sdAB | rabbit | P3 sequence | TTTCTTCATTA | Massive Photonics | 1:500 |
| sdAB | chicken | P7 sequence | TTAATTGAGTA | Massive Photonics | 1:500 |
| sdAB | mouse | F1 sequence | n/a | Massive Photonics | 1:500 |
| sdAB | rabbit | F2 sequence | n/a | Massive Photonics | 1:500 |

**Table 4** Secondary antibodies used for immunostaining.

| Name | Sequence 5'-3' | Dye | Supplier | Concentration (nM) |
|---|---|---|---|---|
| I1 sequence | CTAGATGTAT | Atto647N | Metabion | 0.15 - 0.3 |
| I3 sequence | GTAATGAAGA | Atto655 | Massive Photonics | 0.4 - 0.7 |
| I3 sequence | GTAATGAAGA | Atto680 | Biomers | 0.6 - 1.4 |
| I7 sequence | GTACTCAATT | Atto655 | Massive Photonics | 0.4 - 0.7 |
| F1 sequence | n/a | Atto647N | Massive Photonics | 0.1 |
| F2 sequence | n/a | Atto655 | Massive Photonics | 0.1 |

**Table 5** DNA-PAINT imagers.
